# A Statistical Framework for Robust and Reproducible BioImage Analysis

**DOI:** 10.1101/2025.02.10.637409

**Authors:** David J. Barry, Stefania Marcotti, Lina Gerontogianni, Gavin Kelly

## Abstract

Bioimage analysis is a powerful tool for investigating complex biological processes, but its robustness depends on technical precision and rigorous experimental design. In particular, the use of appropriate controls and experimental repetition is critical for drawing meaningful conclusions. However, both are often used inadequately or overlooked, with “statistical significance” often prioritised, frequently obtained through the misuse or misinterpretation of statistical tests. In this study, we reanalyse publicly available image datasets to highlight the crucial role of robust experimental design in interpreting results. Our findings underscore the importance of focusing on effect sizes and biological relevance over arbitrary statistical thresholds. We also discuss the diminishing returns of increased data collection once statistical stability has been achieved. By refining control usage and emphasising effect sizes, this work aims to enhance the reproducibility and robustness of research findings. We provide open-access code to allow researchers to engage with the dataset, promoting better practices in experimental design and data interpretation.

## Introduction

Bioimage analysis has revolutionised our ability to quantify and interpret complex biological phenomena. However, the robustness of these analyses hinges not only on technical advancements but also on the rigour of experimental design and the correct interpretation of results. Unfortunately, there are many examples in the literature where neither of these criteria are met: experiments are often poorly designed, lack adequate controls, and tend to falsely equate biological and statistical significance. All of these problems stem from a lack of understanding of the variability and measurement noise that is inherent in biological experimentation - a measured difference may appear significant, but this difference could result from day-to-day, sample-to-sample or even cell-to-cell variability, rather than any meaningful biological effect. The biological effect, or effect size, is essentially the difference in the mean value of a measurement between two cell populations, sometimes referred to as the raw effect size.

Despite widespread awareness of their importance, controls are often inadequately implemented or poorly chosen, leading to misinterpretations of experimental results. Effective controls are crucial for distinguishing true biological effects from artifacts introduced by the experimental procedure. However, their role is frequently overshadowed by the pursuit of statistical significance, which is widely acknowledged as a substantial contributor to reproducibility issues (Cobey et al., 2024).

Additionally, despite recognising the importance of repeating experiments to demonstrate reproducible results, researchers often treat individual measurements from within the same sample as “replicates”. Often termed “pseudo-replication” by statisticians, this issue is commonly encountered in bioimage analysis when researchers present measurements from individual cells, drawn from the same sample, as independent data points. These data points are then grouped according to treatment and subjected to statistical tests in pursuit of a “significant” p-value. Applying tests such as ANOVA to these data is inappropriate, as cells from the same sample are necessarily subjected to the same treatment and cannot generally be considered independent. These issues are often exacerbated when non-independent data points from multiple experiments are pooled, artificially inflating sample size and leading to misleadingly low p-values (Lord et al., 2020).

This pursuit of statistical significance can obscure the importance of technically sound experimental design, as small or irrelevant effects may appear significant in the absence of rigorous comparative frameworks. Furthermore, the use of inappropriate or recycled controls undermines the biological validity of findings, leading to conclusions that lack reproducibility. As bioimage data often involve complex and variable biological systems, interpreting results without the proper contextual grounding provided by controls can lead to misleading conclusions.

It is also crucial to address the misconception that a larger sample size always yields more reliable results. Contrary to popular belief, collecting an excessive number of data points to achieve statistical significance is often unnecessary and can lead to a wasteful allocation of resources. The distribution of values within a large population can often be well-represented by a relatively small sample size. While increasing this sample size will result in greater confidence in the obtained statistical descriptors, the law of diminishing returns eventually kicks in. The point at which one should cease collecting data ultimately depends on the level of confidence needed in the statistical description of a particular population, which in turn depends on the size of the effect one is attempting to detect. However, it is important to acknowledge that, even with a well-designed experiment and well-chosen controls, the results can still be ambiguous, and researchers need to be more accepting of this uncertainty in their results.

In this study, we focus on the critical role of experimental controls in bioimage analysis, using publicly available datasets to demonstrate how conclusions can differ significantly based on control selection. Additionally, we highlight that, even with appropriate controls, it is rarely possible to draw conclusions from the results of just one single experiment - repetition is necessary to achieve reproducible results. Our goal is to emphasise the importance of robust experimental design and the importance of control conditions that are not just technically adequate, but biologically meaningful.

While there are many guides on statistics for biologists, such as the excellent ‘Points of Significance’ series in Nature (Krzywinski and Altman, 2013) and the recently published work by Zweifach (2024), they can be perceived as somewhat general, abstract, and difficult to apply to actual experimental results. Pylvänäinen et al. (2025) recently published a practical guide for cell biologists and while they do provide some useful guidance on statistical concepts, their focus was more on data visualisation and exploration. Here, we present analyses of previously published image data, emphasizing that (a) interpreting biological effects is challenging without appropriate controls, (b) when interpreting effect sizes, there is a diminishing return with continued data collection and (c) interpreting the significance of a result is challenging in the absence of repetition. To quote from Fisher and Prance (1974): “*No isolated experiment, however significant in itself, can suffice for the experimental demonstration of any natural phenomenon; for the ‘one chance in a million‘ will undoubtedly occur … In order to assert that a natural phenomenon is experimentally demonstrable we need, not an isolated record, but a reliable method of procedure.*”

We have made the code used for generating the figures in this manuscript openly accessible online, encouraging readers to engage with the data themselves and reproduce the figures using their own datasets. This approach, augmented by actual microscopy data, seeks to make these statistical concepts tangible and directly applicable for cell biologists. It is our hope that this helps to “demystify” statistical analysis, in the context of bioimage analysis, encouraging greater transparency and reproducibility. Additionally, we aim to inspire researchers to embrace ambiguity and uncertainty to a much greater extent.

This study utilises two datasets, IDR0028 (Pascual-Vargas et al., 2017) and IDR0129 (Lawson et al., 2022), freely available from the Image Data Resource (https://idr.openmicroscopy.org; Williams et al. (2017)). The manuscript is divided into approximately three main sections. In the first, we explore how statistical descriptors of a population, such as mean, median and standard deviation, change as more data is collected. We argue that data collection should cease once those descriptors stabilise. We then demonstrate that, within the context of a single experiment, collecting data beyond a certain point yields no further insight into the result. Additionally, we discuss the challenge of interpreting this result in the absence of appropriate controls. Finally, we demonstrate that experimental repeats are necessary in order to draw firm conclusions.

## Results & Discussion

### Overview

In the following we reanalyse subsets of data previously published by Lawson et al. (2022) and Williams et al. (2017). We chose these datasets as they met the following desirable criteria: (1) the images are publicly available via the Image Data Resource (https://idr.openmicroscopy.org), (2) the datasets contain appropriate controls, (3) the data consists of 2D images of cultured cells, labelled with nuclear and actin markers, and are thus relatively easy to analyse, (4) it was possible to extract an intuitive and easy-to-grasp metric quantifying the proportion of a protein in the cell nucleus. Details of all data used is listed in Table 1.

**Table 1.** Overview of the image data used in this study. All images are publicly available from the Image Data Resource (https://idr.openmicroscopy.org).

| Index | IDR Accession Number | Plate | Well | Treatment | Reference |
| --- | --- | --- | --- | --- | --- |
| 1 | idr0139 | 1093711385 | J05 | Untreated | Lawson et al. (2022) |
| 2 | idr0139 | 1093711385 | I19 | Untreated | Lawson et al. (2022) |
| 3 | idr0139 | 1093711385 | G15 | Untreated | Lawson et al. (2022) |
| 4 | idr0139 | 1093711385 | E22 | SN0212398523 | Lawson et al. (2022) |
| 5 | idr0139 | 1093711385 | L08 | Leptomycin B | Lawson et al. (2022) |
| 6 | idr0139 | 1093711385 | L18 | Leptomycin B | Lawson et al. (2022) |
| 7 | idr0139 | 1093711385 | H13 | Leptomycin B | Lawson et al. (2022) |
| 8 | idr0139 | 1093711385 | O02 | DMSO | Lawson et al. (2022) |
| 9 | idr0139 | 1093711385 | B02 | DMSO | Lawson et al. (2022) |
| 10 | idr0139 | 1093711385 | N12 | DMSO | Lawson et al. (2022) |
| 11 | idr0028 | LM2_ONTARGETPlus_1A | I23 | LATS1 RNAi | Pascual-Vargas et al. (2017) |
| 12 | idr0028 | LM2_ONTARGETPlus_1A | J23 | LATS1 RNAi | Pascual-Vargas et al. (2017) |
| 13 | idr0028 | LM2_ONTARGETPlus_1A | M23 | YAP RNAi | Pascual-Vargas et al. (2017) |
| 14 | idr0028 | LM2_ONTARGETPlus_1A | N23 | YAP RNAi | Pascual-Vargas et al. (2017) |
| 15 | idr0028 | LM2_ONTARGETPlus_1A | B06 | Mock Transfection | Pascual-Vargas et al. (2017) |
| 16 | idr0028 | LM2_ONTARGETPlus_2A | C13 | ARAP2 RNAi | Pascual-Vargas et al. (2017) |
| 17 | idr0028 | LM2_ONTARGETPlus_2A | J23 | LATS1 RNAi | Pascual-Vargas et al. (2017) |
| 18 | idr0028 | LM2_ONTARGETPlus_2A | N23 | Mock Transfection | Pascual-Vargas et al. (2017) |
| 19 | idr0028 | LM2_ONTARGETPlus_2A | K16 | YAP RNAi | Pascual-Vargas et al. (2017) |
| 20 | idr0028 | LM2_ONTARGETPlus_2B | C13 | ARAP2 RNAi | Pascual-Vargas et al. (2017) |
| 21 | idr0028 | LM2_ONTARGETPlus_2B | C17 | ARAP2 RNAi | Pascual-Vargas et al. (2017) |
| 22 | idr0028 | LM2_ONTARGETPlus_2B | P13 | Mock Transfection | Pascual-Vargas et al. (2017) |

To analyse the images, we composed a simple and openly available pipeline using CellProfiler (Stirling et al., 2021) to quantify the proportion of a protein of interest that localised to the nuclei (*L_F_*) under different experimental conditions (Fig. 1B). This was achieved by dividing the mean nuclear intensity of the protein of interest (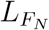) by the mean cellular intensity (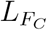):

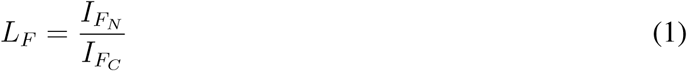

**Fig 1.**
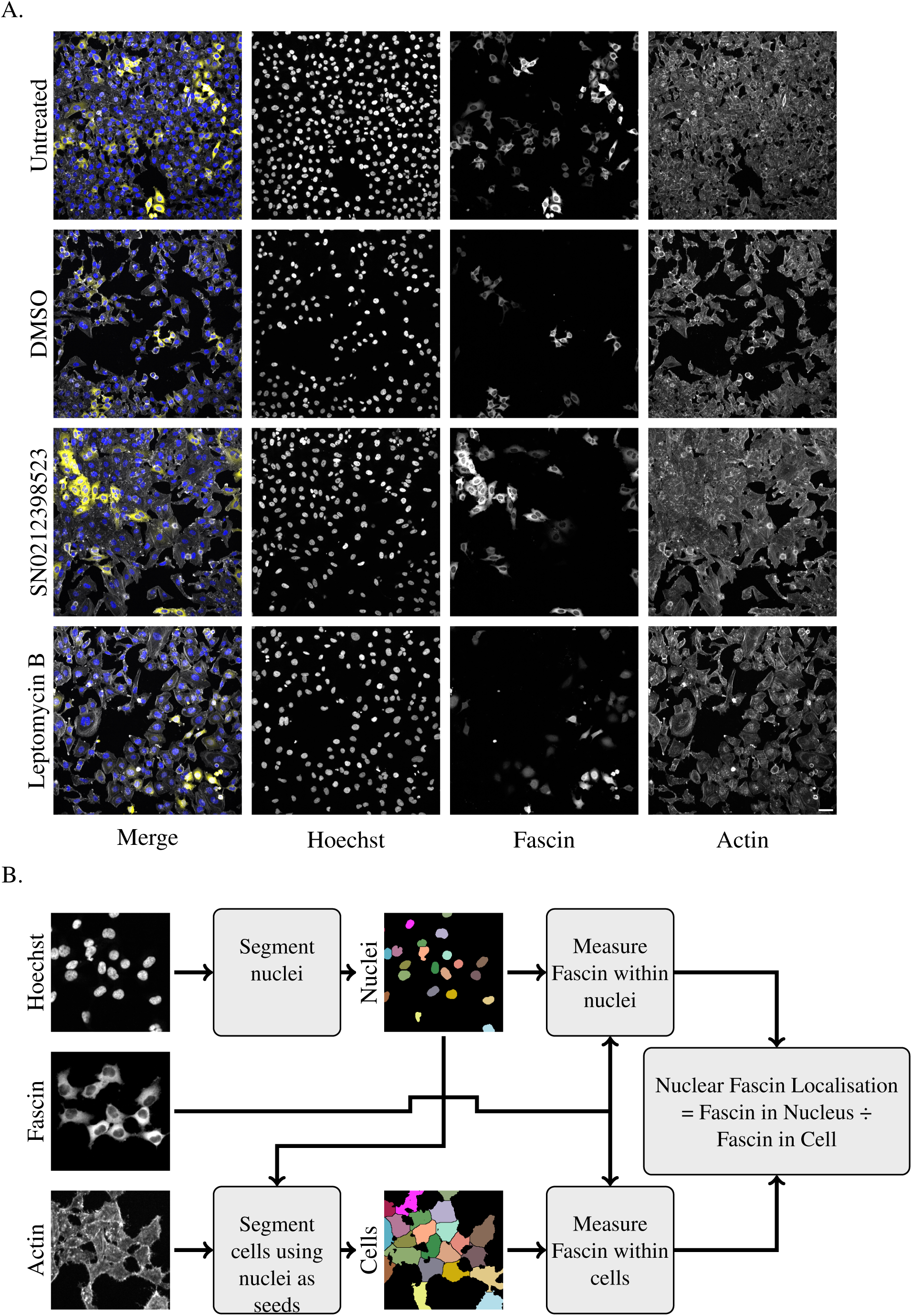
Overview of data and analysis workflow. A. Images of Fascin knockdown (KD) HeLa cells downloaded from the Image Data Resource (accession number IDR0139), originally published by (Lawson et al., 2022). The selected dataset includes a treated sample (pharmacological compound SN0212398523) and three controls (untreated or negative control, DMSO or vehicle control, and Leptomycin B or positive control). The images contain three channels, Hoechst (nuclei), Fascin and actin (cell body). Scale bar 50 *µ*m. B. Schematic illustrating the calculation of relative nuclear Fascin intensity. Fascin localisation is calculated as the mean intensity of Fascin in the nucleus divided by the mean intensity of Fascin in the whole cell. The CellProfiler (Stirling et al., 2021) pipeline used to perform these calculations is openly available in the GitHub repo accompanying this paper (https://github.com/FrancisCrickInstitute/Enhancing-Reproducibility).

### How many cells are needed for a statistically sound result?

Lawson et al. (2022) examined the effects on nuclear localisation of Fascin following treatment with a library of pharmacological compounds (Fig. 1A). The compounds are part of AstraZeneca’s annotated screening library and, as such, their function is not known (https://openinnovation.astrazeneca.com/preclinical-research/target-identification). Here we selected one of these compounds, SN0212398523, for reanalysis. However, the purpose of this work is not to attempt to elucidate any of its biological or clinical roles. The aim here is to illustrate that, in a particular experiment, a result can be obtained with high confidence by analysing a relatively small number of cells.

However, drawing conclusions about biological effects is difficult in the absence of appropriate controls and experimental replication, due to the technical variation inherent to any experiment. For example, even with the use of robotic automation, the concentration of an antibody or dye will be slightly different in different wells. A slight incline in the microscope stage could cause slight differences in the location of the focal plane between wells. Both of these variables will have an influence on the output of an image analysis pipeline, perhaps causing slight differences in the results of cell or nuclear segmentation and the resultant intensity measurements made within those regions.

Lawson et al. (2022) used HeLa cells in which the wild type Fascin was knocked down and replaced with Fascin-mScarlet. We considered three channels in the images, highlighting cell nuclei (Hoescht), Fascin and actin; we used the actin channel as a proxy for the cell body (Fig. 1B). The results of analysing all cells in each of four different wells, corresponding to treatment with the selected compound and three experimental controls, are presented in Table 2. We will begin by considering these results, comparing the test treatment (SN0212398523) to each control one by one. We will then examine how these results change when only a subset of cells from each well is analysed.

**Table 2.** Statistical properties of the data produced by our analysis pipeline. The Index column corresponds to the Index listed in Table 1. The metric being compared, nuclear Fascin localisation, is calculated according to Equation 1

| Index | Treatment | Mean | Median | 25th Percentile | 75th Percentile | Standard Deviation | n<sub>cells</sub> |
| --- | --- | --- | --- | --- | --- | --- | --- |
| 1 | Untreated | 0.4649 | 0.5108 | 0.4177 | 0.5465 | 0.1306 | 1362 |
| 4 | SN0212398523 | 0.4882 | 0.5328 | 0.4479 | 0.5578 | 0.1398 | 844 |
| 5 | Leptomycin B | 0.5791 | 0.5693 | 0.5431 | 0.6262 | 0.1082 | 606 |
| 8 | DMSO | 0.4757 | 0.5241 | 0.4409 | 0.5544 | 0.1308 | 1299 |

## Analysis of all cells in each well

### Treated cells versus negative control

Let us begin by estimating the effect of treating the cells with SN0212398523, relative to untreated cells (negative control). If we were to compare the mean values for these two treatments, it would appear that SN0212398523 causes an increase in Fascin localisation in the nucleus (Table 2). The size of the effect appears small (0.4882 versus 0.4649 - an increase of approximately 2.3%), but we should consider comparisons with appropriate controls before reaching any conclusions.

### Treated cells versus vehicle control

Using untreated cells as a control when the drug is dissolved in DMSO can introduce bias, as, while unlikely, DMSO itself may influence cellular behaviour. A more appropriate comparison would be between drug-treated cells (dissolved in DMSO) and cells treated with DMSO alone. DMSO may introduce a systematic technical variation that could influence the results. The observed effect in SN0212398523-treated cells is likely a combination of technical variation, a biological component due to the active agent, and random variation. Therefore, additional controls are required to conclusively determine any biological significance.

Our results show that treatment with DMSO (vehicle control) leads to an apparent increase in nuclear Fascin localization relative to untreated cells (0.4757 versus 0.4649; Table 2). Furthermore, nuclear Fascin localization in SN0212398523-treated cells was higher than in DMSO-treated cells (0.4882 versus 0.4757; Table 2). This observed effect of DMSO may not be biologically relevant (we would not expect DMSO to have any biological effect on the cells), it is important to note that technical variation can still lead to detectable differences. Researchers must be aware that apparently significant differences can stem from random variation rather than biological effects. Indeed, these differences can be statistically significant, but this does not guarantee biological significance.

### Treated cells versus positive control

Leptomycin B inhibits nuclear export (Lawson et al., 2022), which should result in Fascin accumulating in the nucleus (positive control). Quantifying the proportion of Fascin localised in the nucleus in Leptomycin B-treated cells, we find that the mean value is much larger than all other treatments (Table 2). This helps to put the other measured differences in context. For example, the difference in nuclear Fascin localisation between Leptomycin B-treated cells and SN0212398523 is 0.5791*−*0.4882 = 0.0909, which is several multiples of the difference between SN0212398523-treated and DMSO-treated cells (0.4882 *−* 0.4757 = 0.0125; Table 2).

However, it is important not to over-interpret this result, as it is based on a single experimental replicate. There is a possibility that any of the selected wells may be outliers and not truly representative of the treatment group due to natural variation or experimental error. Without additional replicates to validate the findings, the results may not be generalizable. In other words, the result might not be reproducible, and conclusions drawn from this single experiment may not hold true under slightly different conditions. It is crucial to acknowledge and be aware of this limitation when interpreting results from single experiments. But let us first examine what happens to the above result if we reduce the number of cells analysed in each well.

### How many cells need to be analysed to accurately describe the population in a given well?

Up to this point, we have only considered results derived from an analysis of all the cells in each well, ranging from 606 to 1,362, depending on the treatment (Table 2). We wanted to determine whether similar results could be obtained from an analysis of a subset of cells from each well. Specifically, we wanted to estimated the smallest sample that could be taken from each well to arrive at a result comparable to that in Table 2.

It is possible to ascertain at what point data collection should cease by examining statistical properties as data is being collected. In Figure 2A, we show that the uncertainty in a particular statistical descriptor, inter-quartile range, decays exponentially with increasing sample size. In other words, as more data is collected, the error associated with a particular statistical parameter declines in a highly predictable manner. The inter-quartile range, defined as the difference between the 75th and 25^th^ percentiles (see Table 2), is a commonly used measure of the spread of a dataset. It will be familiar to life scientists as the height of the box in a box plot (see Figure 3A-F, for example).

**Fig 2.**
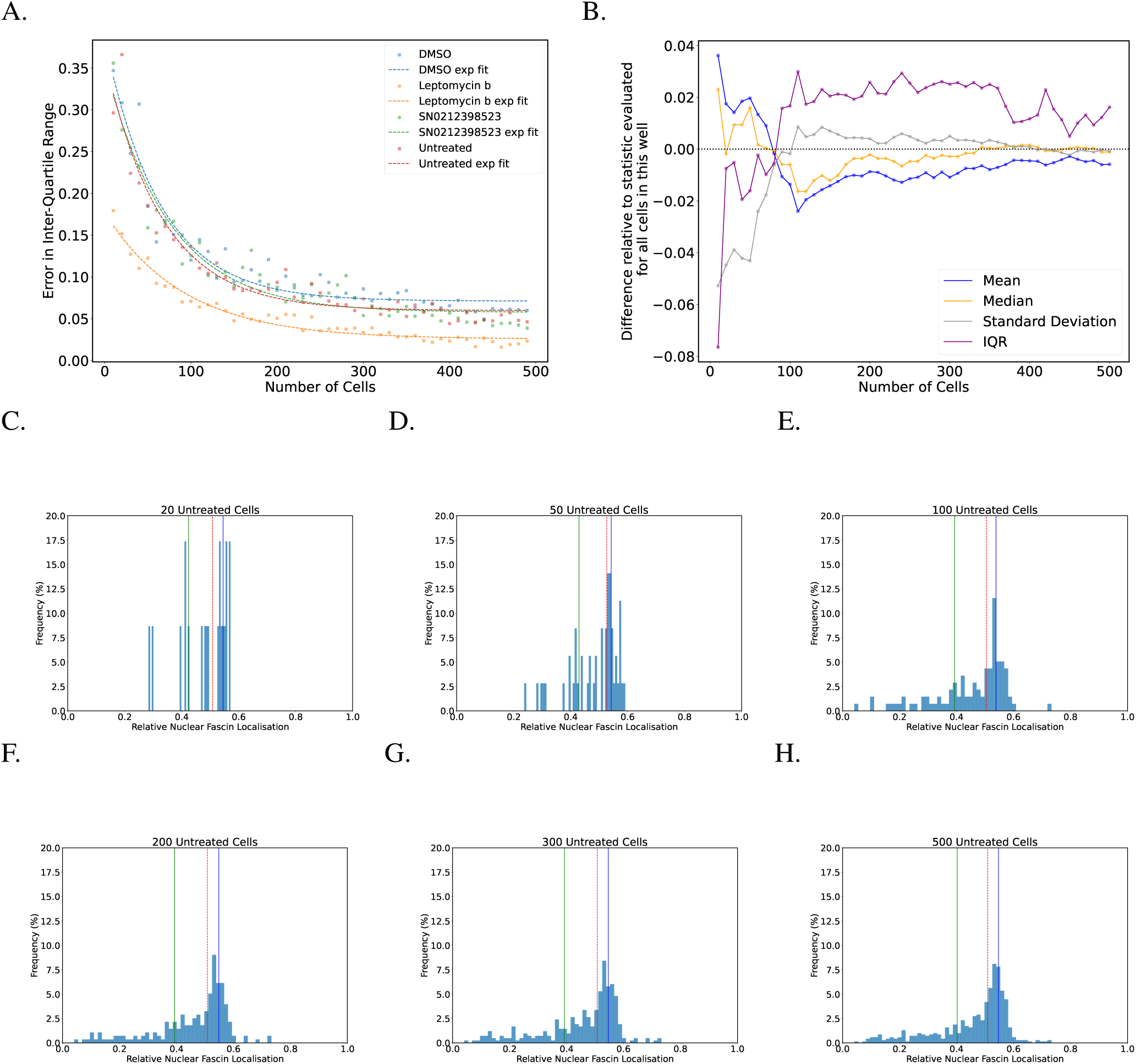
Variation in statistical descriptors stabilises above a certain sample size. A. The difference between the maximum and minimum estimate of the inter-quartile range follows a predictable exponential decay as the number of cells in each sample is increased. Results are based on 100 random cell samples for each sample size. B. The difference between statistical properties of cumulative random cell samples drawn from the population of untreated cells (Table 2) and the properties listed in Table 2 for the entire population. For samples consisting of less than approximately 100 cells, there is substantial variability in the measured statistical properties. However, above a sample size of approximately 200, the statistical properties are relatively stable and tend towards the values listed in Table 2 (the differences tend to zero). C-H. Histograms of the random samples described in ‘B’. Solid vertical green and blue lines depict the width of the inter-quartile range. The dashed vertical red line depicts the median.

**Fig 3.**
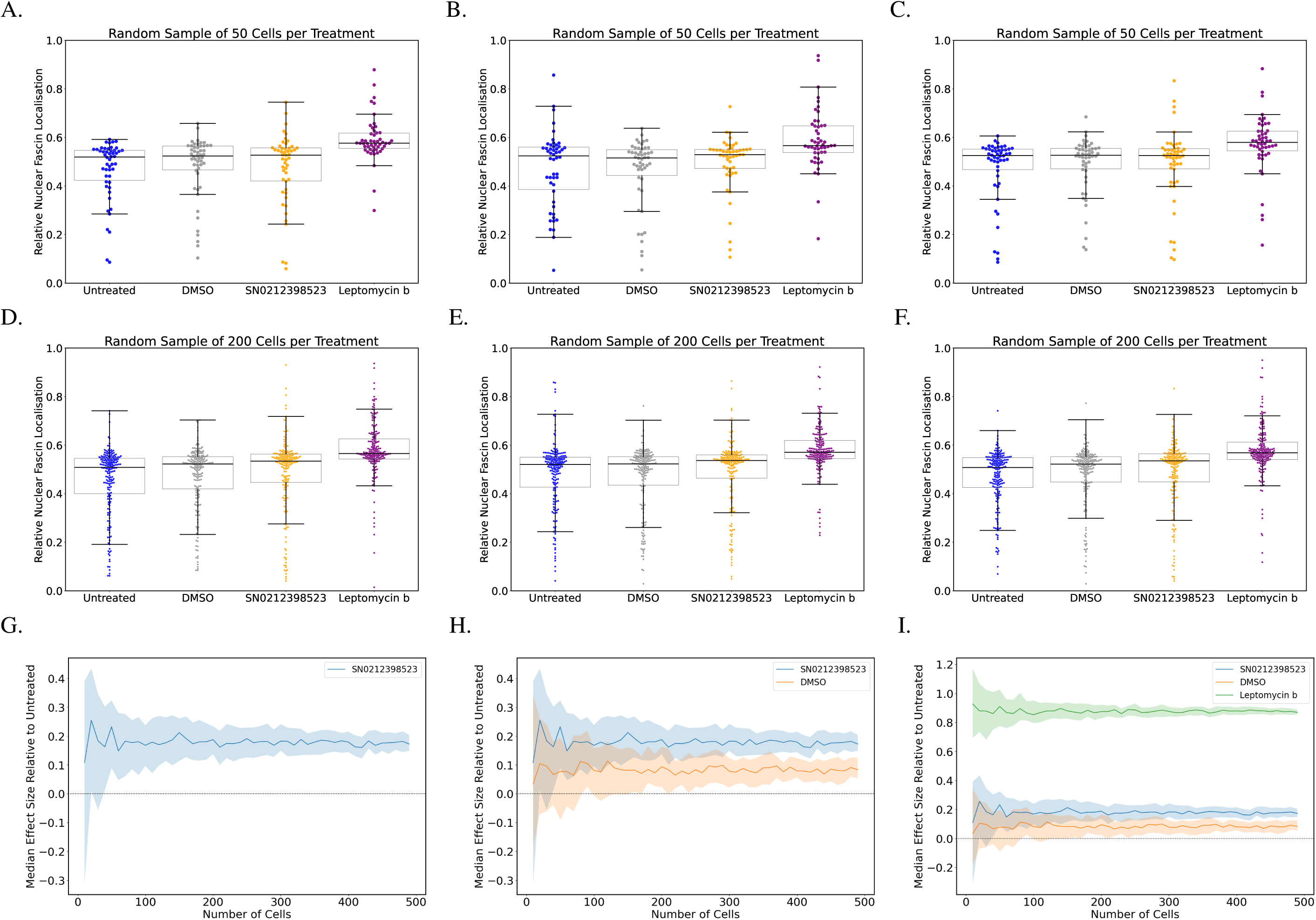
Effect sizes are relatively insensitive to random sampling and sample size. A-F. Three different random samples of 50 (A-C) or 200 (D-F) cells for each treatment-each data point represents a single cell. While the distribution of data points changes in each plot, particularly for the smaller sample size of 50, the overall result is similar in each case. Any random sample of 50 cells or more from these populations will likely show that nuclear Fascin localisation is higher in Leptomycin B-treated cells. G. The influence of random sampling and sample size on the difference in Fascin localisation between treated and untreated cells. For each sample size, random sampling was performed 100 times. The solid line represents the median of those 100 random samples and shaded bands represent the width of the inter-quartile range. Above a sample size of approximately 100, the effect of random sampling on the result is minimal. We can confidently state that nuclear Fascin localisation in treated cells is higher than in untreated cells, but in the absence of controls, interpreting the significance of this difference is difficult. H. A repeat of the bootstrapping analysis in ’G’, but this time including the effect size of measured nuclear Fascin in DMSO-treated cells as a vehicle control when compared to untreated cells. I. One more repeat of the bootstrapping analysis in ’G’ and ’H’, this time including the positive control, Leptomycin B. Effect sizes are calculated according to Equation 2.

This is further illustrated in Figure 2B, which shows how four commonly-used statistical parameters describing the untreated cells data (Table 2) vary as a random sample of the data is increased in size. We observe in our example a threshold effect around a sample size of approximately 250-300 cells, beyond which additional data yields diminishing returns in terms of statistical insight. This can be further seen graphically in Figure2C-H-the distribution of the data for sample sizes between 200 and 500 looks very similar.

This evidence suggests a pragmatic approach to data collection: researchers should aim to gather enough data to reach statistical stability, where the key metrics of interest no longer change significantly with additional sampling. Once this level of statistical confidence is achieved — often visible through a plateau in the variability of statistical parameters — further data collection becomes less informative and may unnecessarily expend resources.

It is important once again at this point to emphasize the difference between increasing the number of data points obtained from a single experiment and increasing the number of independent experiments. In the case of the former, as we discuss above, drawing additional samples (here, cells) from a single experiment will have diminishing returns beyond a certain point-once we have a stable, representative estimate of the mean, for example, additional data may offer little new insight. This differs from collecting independent samples from multiple units, which enhances the generalisability and reliability of the results (Lord et al., 2020). We will revisit this point later in the manuscript.

### How many cells need to be analysed when comparing populations that have undergone different treatments?

The number of cells for each treatment in Table 2 is relatively large and the difference in means is relatively small. However, the data presented in Figure 2 suggests that we should have reasonably high confidence in the result provided by a random sample of approximately 200-300 cells from each treatment group.

We therefore explored the influence of randomly sub-sampling from each well and examining the impact this has on the results (Figure 3A-F). For each of these random samples, the distribution shapes might differ, particularly for the smaller sample size of 50 cells (Figure 3A-C). However, the overall trend in the result is virtually identical-for example, the centreline (median) of the box plots representing Leptomycin B-treated cells is consistently higher than all other treatments.

We further investigated this relationship between sample size (number of cells), random sampling, and effect size. We randomly sampled (with replacement) from the different populations in Table 2 and once again measured the proportion of Fascin localised in the nucleus across all treatment conditions, using sample sizes ranging from 10 up to 500. We repeated this 100 times for each sample size to produce distributions of 100 mean values for each sample size (Figure 3G-I).

Let us first examine the influence of sample size and random sampling on the difference between Fascin localisation in untreated cells and cells treated with SN0212398523 (Figure 3G). For small sample sizes (*<* 50), there is a large amount of uncertainty in measured effect size (the difference in mean nuclear Fascin localisation between the two populations). But, above a sample size of approximately 50, the effect of random sampling is minimal and the result changes little. We can reasonably conclude that, in this particular experiment, for any sample of approximately 50 or more cells, the measured nuclear Fascin localisation is higher in SN0212398523-treated cells than in untreated cells. Adding more cells to the analysis will not alter this conclusion. However, it is not clear how biologically significant this difference is.

We repeated this analysis, but this time included the vehicle control, DMSO-treated cells (Figure 3H). Again, we can see that, above a certain threshold (approximately 250-300 cells per sample) there is little overlap between the inter-quartile ranges and we can reasonably conclude that, according to our analysis pipeline, nuclear Fascin localisation is higher in SN0212398523-treated cells than it is in DMSO-treated cells in this experiment. But the fact that Figure 3H also shows that nuclear Fascin localisation is higher in DMSO-treated cells than it is in untreated cells should cause us to question the biological significance of any measured effect of SN0212398523, because we know that DMSO almost certainly does not promote nuclear localisation of Fascin. At this point, it is difficult to distinguish between technical variability (which likely accounts for the difference between untreated and DMSO-treated cells) and potential biological effects of SN0212398523. In other words, the higher level of nuclear Fascin we have measured in the presence of SN0212398523 might be a real biological effect, or it might just be a technical artifact, caused by random well-to-well variability. Therefore, additional controls are required to aid our interpreation.

Repeating the same analysis one more time with a positive control puts these differences into context (Figure 3I). There are two important points to highlight from these results. The first is that, while there is a measured increase in nuclear Fascin localisation in DMSO-and SN0212398523-treated cells, it does not seem biologically significant, relative to the increase in localisation measured in Leptomycin B-treated cells. The second is that, even for small sample sizes (*<* 50 cells), this result is apparent. Analysing more cells only serves to refine the measured differences (that is, the uncertainty in the estimates of the means is reduced)-it does not alter the overall conclusion. However, in this particular experiment, the uncertainty in the effect of Leptomycin B (the width of the shaded area of the plot in Figure 3I) is small relative to the effect size (the solid lines in Figure 3I). In other scenarios, in which effect sizes are smaller and the associated uncertainty greater, more data collection may be necessary in order to draw conclusions from the results.

### Is the relationship between sample size and effect size reproducible across multiple experiments?

Up to this point, we have only considered data from one single “experiment”, or one replicate. To test the influence of random sampling and sample size when comparing multiple experiments, we analysed additional data from Lawson et al. (2022) (Figure 2). We found that increasing the number of cells analysed and averaged from 50 to 200 had a small influence on the result (Figure 2A & B). The distribution of data points changes slightly, but the overall conclusion does not change-Leptomycin B has a far larger effect on nuclear Fascin localisation than SN0212398523. Consistent with other data presented up to this point, we found that increasing the number of cells in each sample beyond 200 had little effect-we obtained an almost identical result when 500 cells were randomly sampled (Figure 2C). One could employ an ANOVA test at this point to add additional weight to the result. But given that all box and whiskers plots overlap, except for those associated with Leptomycin B, no statistical test should be necessary to conclude that Leptomycin B has a far more significant biological effect than any other treatment.

There is an important point to be made at this juncture regarding the misuse or misinterpretation of p-values. It is not unusual for data similar to that presented in Table 2 to be presented as a swarm plot, annotated with p-values, where each cell is treated as an independent data point (Figure 1). For examples, see Cai et al. (2024), Figure 1 D & E or Buglak et al. (2024), Figure 1 G-I. Without getting into a detailed discussion of what constitutes a technical or biological replicate, it is not unreasonable to state that different cells within the same dish/sample all subjected to the same treatment are *not* replicates (Zweifach, 2024)-they constitute one single experiment. As such, subjecting such data to statistical tests makes little sense. Regardless, blindly relying on the p-values presented in Supplemental Figure 1 might lead us to conclude that SN0212398523 has a highly significant influence on nuclear Fascin localisation. However, all of the data we have presented up to this point suggests that this is not the case.

However, a major limitation of the dataset we have relied on thus far is that there is only one single well, or replicate, for each compound tested-very often a feature of preliminary screens. Hence, in Supplemental Figure 2, we are comparing one well for SN0212398523 with three for the untreated, DMSO- and Leptomycin B-treated populations, which is not ideal. To address this shortcoming, and also illustrate the robustness of our approach, we now apply the same analysis to a second publicly-available dataset.

### Applying our statistical framework to new data

We reanalysed a subset of the data previously published by Pascual-Vargas et al. (2017) (Fig. 4). We chose this particular dataset as, similar to the data published by Lawson et al. (2022), the images are publicly available via the Image Data Resource. The nature of the analysis we conducted is almost identical to that used for the analysis of the data from Lawson et al. (2022), as the images from Pascual-Vargas et al. (2017) also consisted of 2D images of cultured cells, labelled with nuclear and actin markers. We considered the relative localisation of a protein, YAP/TAZ, to the nucleus, which is again intuitive and should easily be grasped by life scientists.

**Fig 4.**
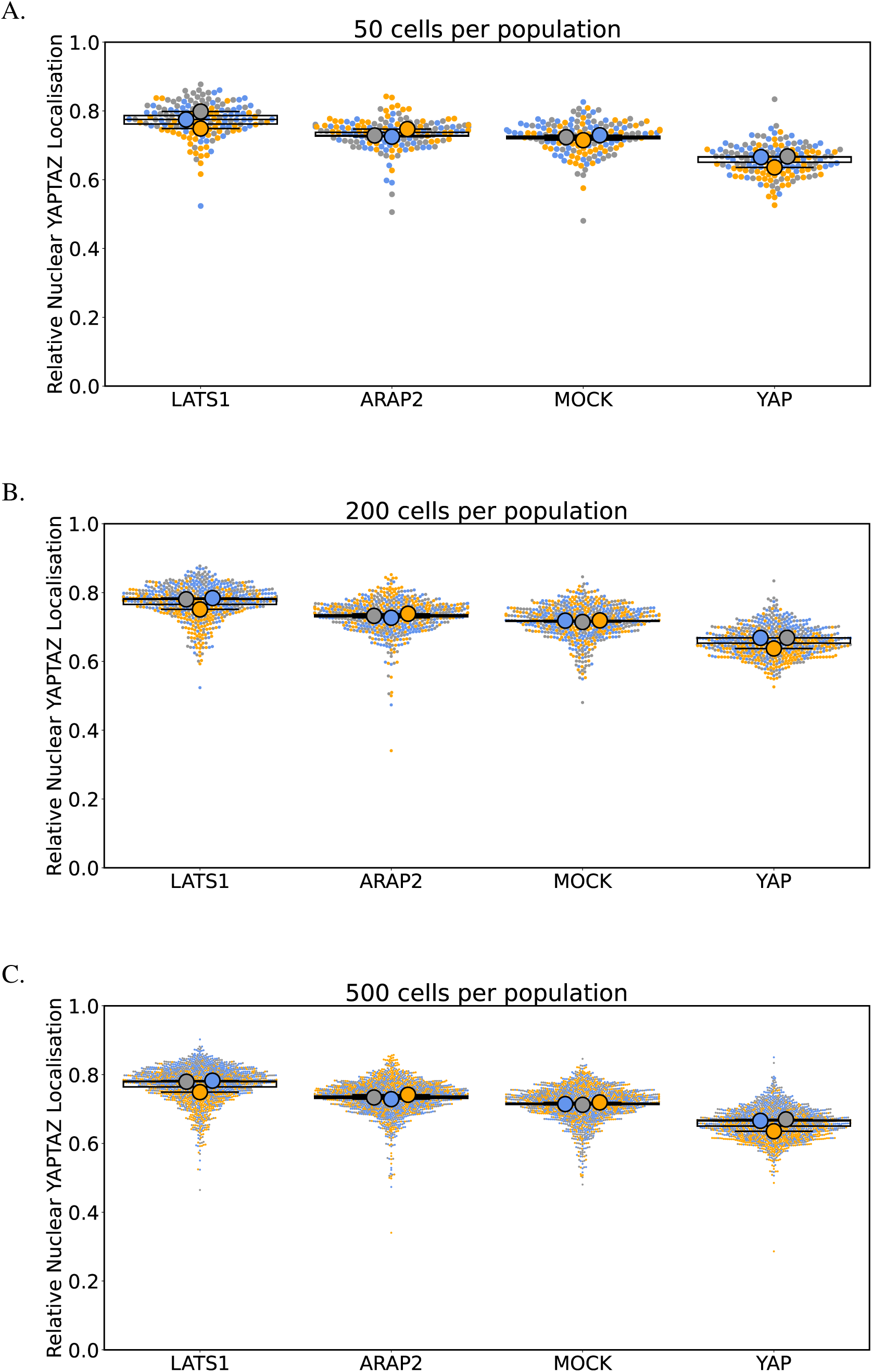
Effect size is relatively insensitive to random sampling when comparing multiple experiments. A-C. Nuclear YAP/TAZ localisation averaged across samples of cells subjected to the indicated siRNA knockdowns. Each large dot represents the average of 50 (A), 200 (B) or 500 (C) randomly selected cells from three different wells. Each small dot represents a single cell. Dots with the same colour correspond to a particular experimental replicate. The source of the data for each replicate are listed in Table 1-all data comes from the same RNAi screen, but wells were selected from three different multi-well plates to simulate experimental repetition.

Pascual-Vargas et al. (2017) conducted an RNAi screen to isolate Rho Guanine Nucleotide Exchange Factors (RhoGEFs) and Rho GTPase-activating proteins (RhoGAPs) that contribute to the morphogenesis of LM2 and MDA-MB-231 cell lines. Here we selected one of these RNAis, against ARAP2, for reanalysis, comparing to control RNAis that promote high (LATS1) or low (YAP) localisation of YAP/TAZ to the cell nucleus and a neutral control (mock transfection). We considered three channels in the images, highlighting cell nuclei (Hoescht), YAP/TAZ and actin, once again using the actin channel as a proxy for the cell body. We used the same CellProfiler pipeline to quantify the proportion of YAP/TAZ that localised to the nuclei of LM2 cells under different experimental conditions.

### Increased data collection again yields diminishing returns

We begin by demonstrating that the general principles outlined in Figure 2 for the analysis of data from Lawson et al. (2022) also hold true for that of Pascual-Vargas et al. (2017) (Supplemental Figure 3). We show again that the uncertainty in the inter-quartile range decays exponentially with increasing sample size (Supplemental Figure 3). We go on to show how the mean, median, inter-quartile range and standard deviation describing the nuclear YAP/TAZ localisation in mock transfected cells vary as a random sample of the data is increased in size. We again observe a dataset-specific threshold effect, this time around a sample size of approximately 200 cells, beyond which additional data yields diminishing returns in terms of statistical insight. We further emphasise this in Supplementary Figure3C-H-the distribution of the data for sample sizes between 200 and 500 looks very similar. But we again emphasise the difference between increasing the number of data points obtained from a single experiment and increasing the number of independent experiments (the latter being explored below).

Collecting additional data points from a single experiment has limited value (Supplemental Figure 3), but collecting independent samples from multiple experiments enhances the generalisability and reliability of the results (Lord et al., 2020).

### Differences in treatment groups can be ascertained from small sample sizes

As we did for the data from Lawson et al. (2022), we next explored the influence of randomly sub-sampling on our analysis of the data from Pascual-Vargas et al. (2017) (Supplemental Figure 4A-F). As before, the overall trend in the result is virtually identical for different sample sizes and the differences in measured effect are similar. Exploring further, we again randomly sampled (with replacement) the data from Pascual-Vargas et al. (2017) and measured the proportion of YAP/TAZ localised in the nucleus across all treatment conditions, using sample sizes ranging from 10 up to 500. We repeated this 100 times for each sample size to produce distributions of 100 mean values (Supplemental Figure 4G-I). We see that, above a sample size of approximately 50-100, the effect of random sampling is minimal and the result changes little. We can reasonably conclude that, in this particular experiment, for any sample of approximately 50 or more cells, the nuclear YAP/TAZ localisation will be higher in ARAP2 RNAi-treated cells than in mock transfected cells (Supplemental Figure 4G). We can also conclude that while ARAP2 RNAi-treated cells may exhibit considerably higher nuclear YAP/TAZ localisation than our negative control (YAP), the effect is not quite as strong as for the positive control (LATS1). This result is evident having analysed 50-100 cells-adding further data-points yields no additional insight. However, it bears repeating that this is the result of one single experiment and we cannot reach any firm conclusions until we examine the results of replicates.

### Relationship between sample size and effect size is reproducible across multiple experiments

To test the influence of random sampling and sample size when comparing multiple experiments, we analysed additional data from Pascual-Vargas et al. (2017) (Figure 4), including two more wells per treatment drawn from multiple plates. We found that increasing the number of cells analysed and averaged from 50 to 200 had almost no influence on the result (Figure 4A & B). The distribution of data points changes slightly, but the overall conclusion does not change-LATS1 RNAi has a larger effect on nuclear YAP/TAZ localisation than ARAP2. Consistent with other data presented up to this point, we found that increasing the number of cells in each sample beyond 200 had little effect-we obtained an almost identical result when 500 cells were randomly sampled (Figure 4C).

### A Framework for Robust Quantitative BioImaging Experimental Design

Our analysis highlights the limitations of over-interpreting small treatment effects, especially without appropriate controls and experimental repeats. Biological significance can only be conclusively inferred when effect sizes are measured against controls that deliver biologically meaningful and measurable effects. Relying solely on arbitrary statistical thresholds, such as p-values, to distinguish between significant and insignificant results can lead to misconceptions about the biological importance of findings. We propose several strategies to help researchers develop a more nuanced understanding of statistical results and to encourage practices that prioritise biological relevance over arbitrary statistical thresholds when designing imaging experiments (Box 1).

Ideally, consult a statistician from the initial stages of research design. Early collaboration helps select appropriate statistical tests, plan effective experiments, and avoid common pitfalls in p-value interpretation. However, many researchers do not have access to such expertise and much of what follows is aimed at researchers who must go it alone.

## 1.0 Perform a Pilot Study

Designing a quantitative imaging experiment for the first time often involves unknowns. Starting with a pilot experiment to obtain preliminary data and establish criteria for larger experiments is sensible. It is of utmost importance that the analysis of the image data is considered from the outset, given equal importance alongside sample preparation and image acquisition. Fully interpreting the resultant data to inform further iterations is essential (Senft et al., 2023).

### 1.1 Determine how many cells are needed per treatment

The number of cells needed for a representative sample depends on the variability in the measured property. Estimating this variability can be challenging, especially with novel cell lines, compounds, and imaging systems common in biological research.

However, the code we provide accompanying this manuscript can be used to arrive at an estimate of appropriate sample size. Acquire some images of some cells, analyse them and quantify the metric of interest, then analyse the statistical properties of the population to see how they change as the number of cells is increased. An estimate for a good samples size can be obtained by observing when the statistical descriptors of the distribution stabilise around a constant value and range.

##### Box 1: Statistical Framework for Robust and Reproducible BioImage Analysis.

1. **Outline Experiment**

- Define the biological question and expected effect size.
- Determine the metric to measure.
2. **Establish Controls**

- Determine the controls to use.
- Prepare individual dishes/wells containing untreated, neutral, positive, and negative controls.
- Determine the number of cells required per dish/well to achieve statistical stability.
- Is the effect of controls detectable?
3. **Establish Reproducibility**

- Repeat the pilot experiment *n* times, where *n* is *>*= 3
- Generate a superplot (Lord et al., 2020). Is the effect detectable reproducibly?
- If not, modify the pilot experiment.
4. **Test Conditions**

- Introduce test condition(s)
- Repeat thr experiment *n* times, where *n* is *>*= 3
- Generate a superplot (Lord et al., 2020). The effect of treatment(s), relative to controls, should be apparent
- Interpret findings by focusing on effect size and biological relevance, comparing observed effects to controls.
5. **Transparent Reporting & Data Sharing**

- Publish image data on the BioImage Archive (Hartley et al., 2022).
- Publish code and pipelines on Github.
- License appropriately.
- Adhere to FAIR principles (Findable, Accessible, Interoperable, Reusable) (Wilkinson et al., 2016).

### 1.2 Estimate the size of the effect to be detected

Repeat step 1.1 for different experimental conditions and determine whether an effect can be detected when comparing treatment versus controls. If not, it may be that additional data is required (return to step 1.1 above), or it may be that there is no measurable effect and a redesign of the experiment might be required.

## 2.0 What controls should I use?

Experiments often default to the simplest control: comparing treated samples against untreated ones. However, this binary approach can overlook nuanced biological responses and fail to account for variables like treatment process effects or batch effects. Moreover, the practice of reusing control samples across different experimental repeats—a shortcut that compromises the integrity of the data—highlights the need for a more rigorous approach to control selection.

### 2.1 What if there are no appropriate positive/negative controls?

Depending on the situation, non-biological controls might be appropriate. For example, if measuring co-localisation of punctate proteins, multi-labelled fluorescent beads may be an appropriate positive control and comparing the signal in one channel with noise in another might constitute an appropriate negative control.

## 3.0 How many replicates to I need?

Repeat your pilot experiment. Are the results reproducible? Produce a superplot (Lord et al., 2020). Is there a clear pattern or trend? If yes, scale up your experiment and attempt to reproduce the result. If the results vary considerably, redesigning the experiment may be necessary.

### 3.1 What constitutes a replicate?

Ideally, every aspect of an experiment should be independent from all others. However, there are circumstances in which this may be impractical (for example, perhaps access to a microscope is limited). In such cases, independence must be maintained so far as is possible. But at the absolute minimum, in order for two different cell populations to be considered “independent“, they *must* be in different wells or dishes, assuming all wells in a single well or dish are being subjected to the same treatment.

## 4.0 Transparent Reporting and Data Sharing

All image data used in this work were freely shared online, allowing us to reuse it. We have made all the code used to generate the figures in this manuscript freely available. We encourage readers to publish their work in a similar open and accessible manner. We strongly recommend that all bioimage data be published on the BioImage Archive (Hartley et al., 2022) and that all image analysis pipelines and associated code be published in an online repository such as GitHub or similar. We suggest licensing your material appropriately, such as using the Creative Commons Attribution 4.0 International Public License (CCB). Further guidelines on publishing image analysis pipelines have been published elsewhere (Schmied et al., 2024; Miura and Nørrelykke, 2021). As far as possible, all data should meet the FAIR principles (Findable, Accessible, Interoperable, Reusable) for scientific data management and stewardship (Wilkinson et al., 2016).

## Conclusion

Our findings emphasise the importance of using appropriate controls and focusing on effect sizes rather than, for example, p-values. By considering biological relevance over arbitrary thresholds of significance, researchers can derive more robust and reproducible conclusions. These recommendations aim to guide future bioimage analysis toward more meaningful interpretations and efficient experimental designs.

We caution against solely relying on statistical tests for interpreting bioimage data. Collaborating with statisticians for data analysis is highly recommended. More importantly, we stress the use of appropriate controls to discern biological meaningfulness. Although this might introduce greater uncertainty in research outcomes, it is a necessary step towards reducing irreproducible “highly significant” results, thereby advancing the credibility and robustness of scientific research.

## Materials & Methods

### Image Data

The image data used in this study, first published by Lawson et al. (2022) and Pascual-Vargas et al. (2017), is freely available on the Image Data Resource (https://idr.openmicroscopy.org) under accession numbers IDR0028 and IDR0139. A subset of each dataset was used, the details of which are listed in Table 1.

### Image Analysis

All image analysis was performed with a custom image analysis pipeline implemented in CellProfiler (Stirling et al., 2021), which is freely available on Github, along with all other code associated with this study (https://github.com/FrancisCrickInstitute/Enhancing-Reproducibility). Briefly, the Hoechst and actin channel of each image were smoothed using Gaussian filtering. Nuclei were then identified in the smoothed Hoechst channel and cell bodies in the smoothed actin channel using minimum cross-entropy thresholding. Measurements of the protein of interest (either Fascin or YAP/TAZ) were then made within these segmented objects. To obtain the percentage of nuclear protein used as quantitative output, the signal intensity in the nuclear region was divided by the signal intensity in the whole cell.

### Data Analysis

All data analysis was performed in Python using the numpy (Harris et al., 2020), pandas (McKinney, 2010), scikit-posthocs (Terpilowski, 2019) and scipy (Virtanen et al., 2020) packages. Plots were generated using matplotlib (Hunter, 2007) and seaborn (Waskom, 2021). Companion Jupyter notebooks, inclusive of instructions on how to set them up locally using Python environments, is available on GitHub (https://github.com/FrancisCrickInstitute/Enhancing-Reproducibility).

Individual data points in swarm plots represent individual cells, unless otherwise stated. Box-and-whiskers plots show median (central line), inter-quartile range (box edges) and 1.5 times the inter-quartile range (whiskers). Effect size (*d*) is calculated according to Cohen (2013) as:

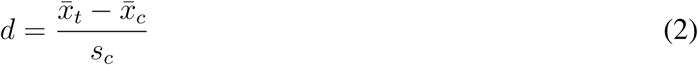

where *x̄_t_* and *x̄_c_* represent the mean of the treated and control cell samples and *s_c_* is the standard deviation of the control cell sample.

## Funding

This work was supported by the Francis Crick Institute, which receives its core funding from Cancer Research UK (CC1069, CC1107), the UK Medical Research Council (CC1069, CC1107) and the Wellcome Trust (CC1069, CC1107). SM has received funding from the Biotechnology and Biological Sciences Research Council (BBSRC, grant number BB/V006169/1).

## Code Availability

All code used to generate the figures in this manuscript is available online (https://github.com/FrancisCrickInstitute/Enhancing-Reproducibility).

## Acknowledgements

The authors would like to thank Maddy Parsons for constructive feedback on the manuscript.

---

**Supplemental Figure 1.**
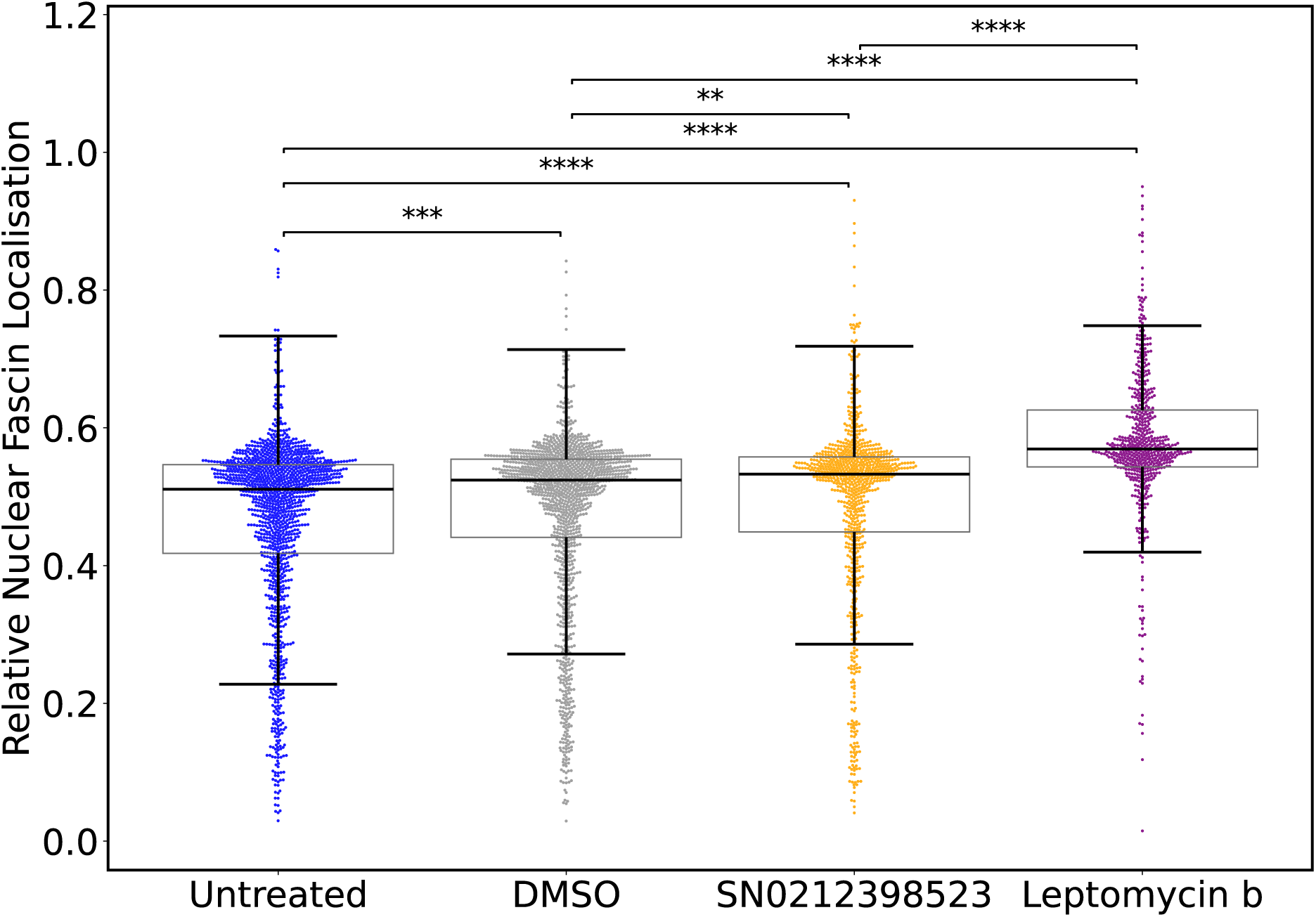
Graphical representation of the data shown in Table 2. Each data point represents a single cell from one of four different wells (corresponding to each of the four different treatments shown) on the same multi-well plate (see Table 1). The plot has been annotated with representations of significant differences, but these differences have been calculated using a Kruskal-Wallis test followed by Dunn’s multiple comparisons, which is inappropriate in this case, as the data points (cells in the same wells) are not independent. We highlight this here as it is not uncommon to see statistical tests misused in this way in the literature-for example, see Cai et al. (2024) or Buglak et al. (2024). Because there are so many data points in each distribution, the p-values obtained are extremely small, potentially misleading an observer into believing that there are highly significant differences between each population. **: *p <* 0.01. ***: *p <* 0.001. ****: *p <* 0.0001.

**Supplemental Figure 2.**
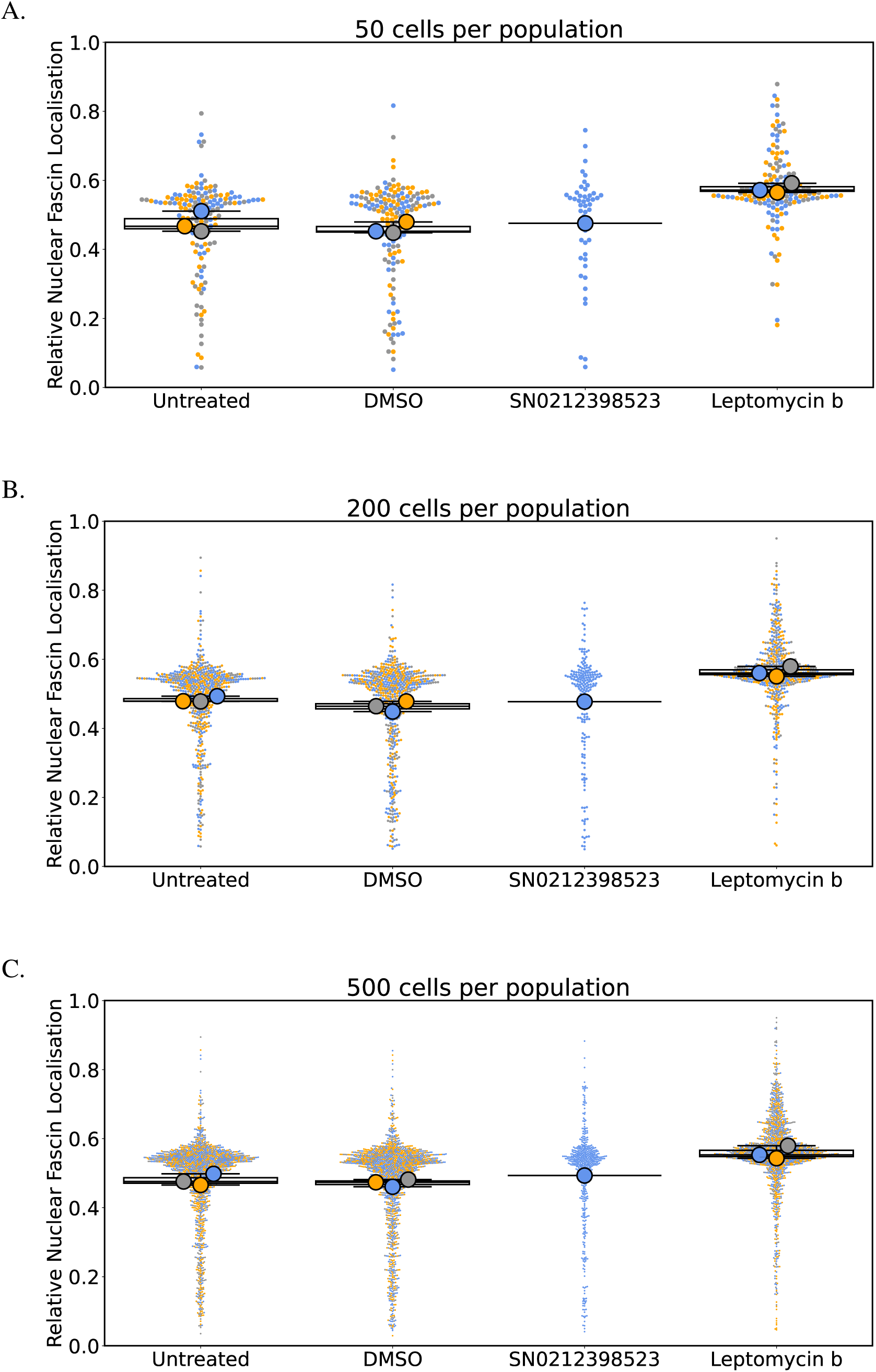
Effect size is relatively insensitive to random sampling when comparing multiple experiments. A-C. Nuclear Fascin localisation averaged across samples of cells subjected to the indicated treatments. Each large dot represents a technical replicate and is the average of 50 (A), 200 (B) or 500 (C) randomly selected cells within a well. Each small dot represents a single cell. Colours correspond to a particular experimental replicate.

**Supplemental Figure 3.**
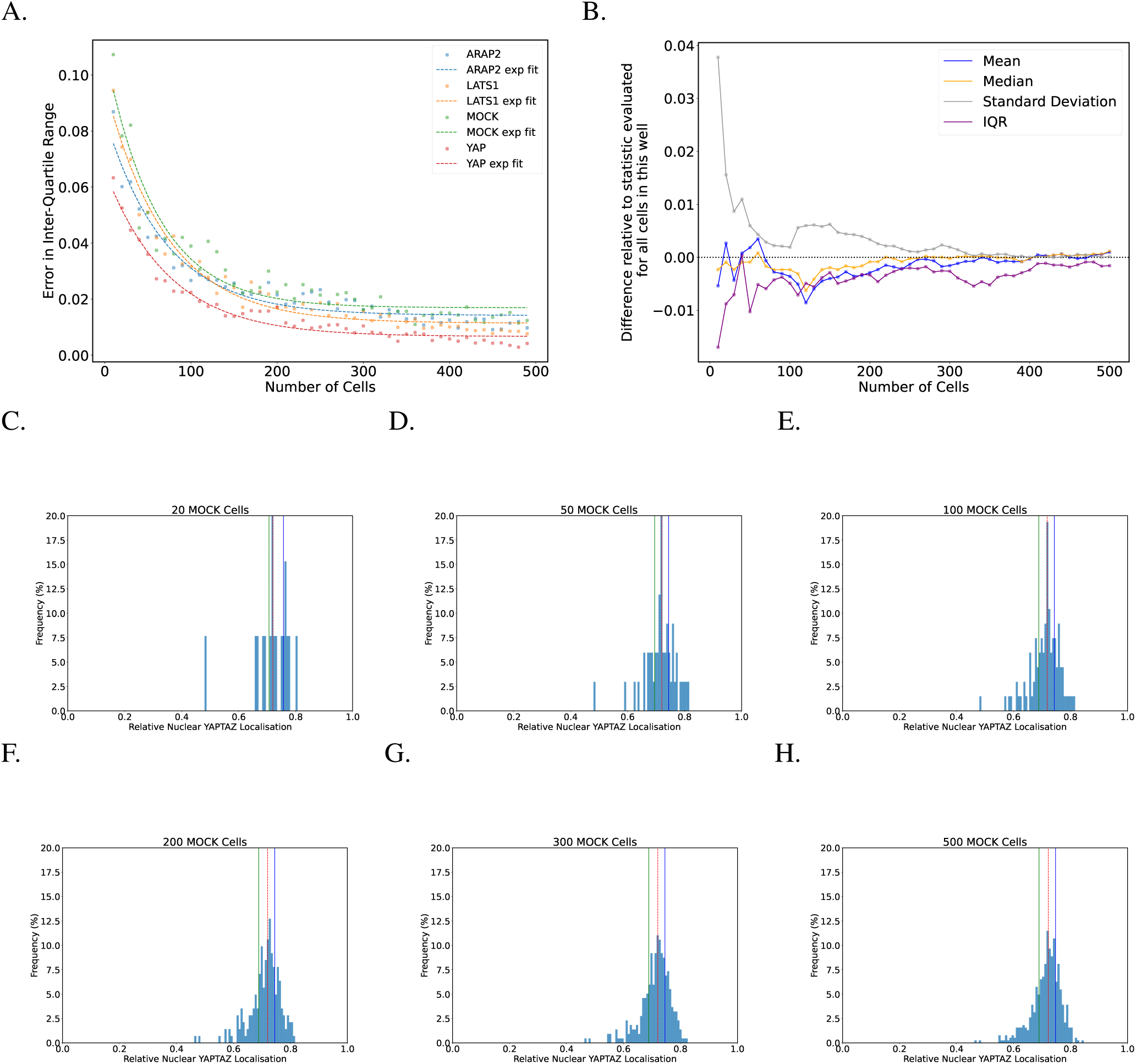
Variation in statistical descriptors stabilises above a certain sample size. A. The difference between the maximum and minimum estimate of the inter-quartile range follows a predictable exponential decay as the number of cells in each sample is increased. Results are based on 100 random cell samples for each sample size. B. The difference between statistical properties of cumulative random cell samples drawn from the population of MOCK-transfected cells (Table 1) and the statistical properties calculated for the entire population. For samples consisting of less than approximately 100 cells, there is substantial variability in the measured statistical properties. However, above a sample size of approximately 200, the statistical properties are relatively stable and the differences between sample statistics and population statistics tend towards zero. C-H. Histograms of the random samples described in ‘B’. Solid vertical green and blue lines depict the width of the inter-quartile range. The dashed vertical red line depicts the median.

**Supplemental Figure 4.**
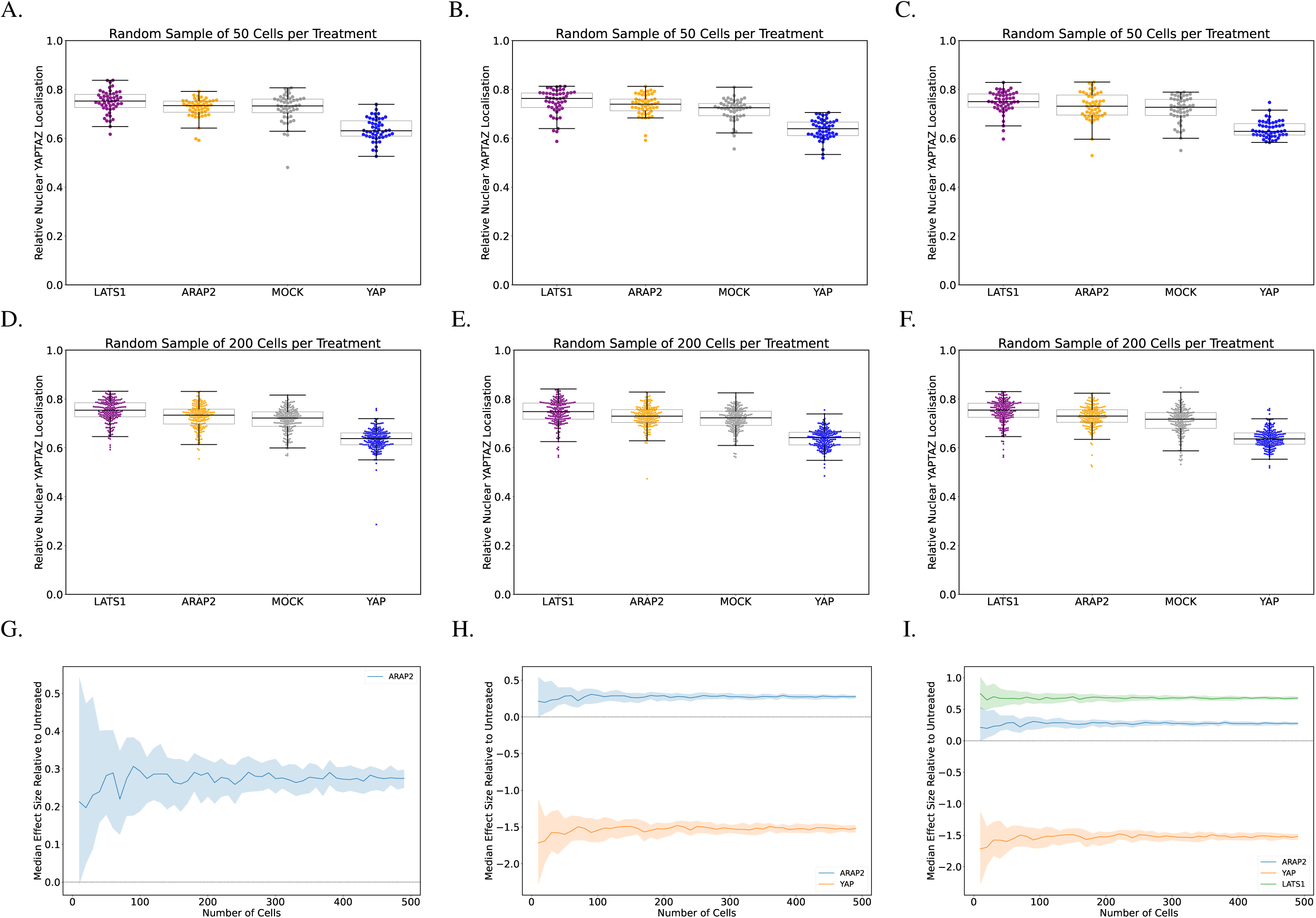
Effect sizes are relatively insensitive to random sampling and sample size. A-F. Three different random samples of 50 (A-C) or 200 (D-F) cells for each RNAi treatment-each data point represents a single cell. While the distribution of data points changes in each plot, particularly for the smaller sample size of 50, the overall result is similar in each case. Any random sample of 50 cells or more from these populations will likely show that nuclear YAP/TAZ localisation is higher in LATS1 RNAI-treated cells and lower in YAP RNAi-treated cells. G. The influence of random sampling and sample size on the difference in YAP/TAZ localisation between RNAi-treated and MOCK-transfected cells. For each sample size, random sampling was performed 100 times. The solid line represents the median of those 100 random samples and shaded bands represent the width of the inter-quartile range. Above a sample size of approximately 50, the effect of random sampling on the result is minimal. We can confidently state that nuclear YAP/TAZ localisation in ARAP2 RNAi-treated cells is higher than in MOCK-transfected cells, but in the absence of controls, interpreting the significance of this difference is difficult. H. A repeat of the bootstrapping analysis in ’G’, but this time including the effect size of measured nuclear YAP/TAZ in YAP RNAi-treated cells as a negative control when compared to MOCK-transfected cells. I. One more repeat of the bootstrapping analysis in ’G’ and ’H’, this time including the positive control, LATS1 RNAi. Effect sizes are calculated according to Equation 2.

## Notes

https://github.com/FrancisCrickInstitute/Enhancing-Reproducibility

### Competing Interest Statement

The authors have declared no competing interest.

### Summary of Updates

This version of the manuscript includes the application of our framework to a second publicly available dataset from the Image Data Resource. Additionally, we have reformatted our recommendations to align more closely with the outline of a framework for others to follow in their own work.

## References

1. Cobey, K. D., Ebrahimzadeh, S., Page, M. J., Thibault, R. T., Nguyen, P.-Y., Abu-Dalfa, F., and Moher, D. (2024) Biomedical researchers’ perspectives on the reproducibility of research. PLOS Biology 22, 1–15.

2. Lord, S. J., Velle, K. B., Mullins, R. D., and Fritz-Laylin, L. K. (2020) SuperPlots: Communicating reproducibility and variability in cell biology. Journal of Cell Biology 219, e202001064.

3. Krzywinski, M., and Altman, N. (2013) Importance of being uncertain. Nature Methods 10, 809–810.

4. Zweifach, A. (2024) Determining how many cells to average for statistical testing of microscopy experiments. Journal of Cell Biology 223, e202401074.

5. Pylvänäinen, J. W., Grobe, H., and Jacquemet, G. (2025) Practical considerations for data exploration in quantitative cell biology. Journal of Cell Science 138, jcs263801.

6. Fisher, R., and Prance, G. The Design of Experiments; Hafner Press, 1974.

7. Pascual-Vargas, P., Cooper, S., Sero, J., Bousgouni, V., Arias-Garcia, M., and Bakal, C. (2017) RNAi screens for Rho GTPase regulators of cell shape and YAP/TAZ localisation in triple negative breast cancer. Scientific Data 4, 170018.

8. Lawson, C. D., Peel, S., Jayo, A., Corrigan, A., Iyer, P., Dalrymple, M. B., Marsh, R. J., Cox, S., Audenhove, I. V., Gettemans, J., and Parsons, M. (2022) Nuclear fascin regulates cancer cell survival. eLife 11, e79283.

9. Williams, E., Moore, J., Li, S. W., Rustici, G., Tarkowska, A., Chessel, A., Leo, S., Antal, B., Ferguson, R. K., Sarkans, U., Brazma, A., Salas, R. E. C., and Swedlow, J. R. (2017) Image Data Resource: a bioimage data integration and publication platform. Nature Methods 14, 775–781.

10. Stirling, D. R., Swain-Bowden, M. J., Lucas, A. M., Carpenter, A. E., Cimini, B. A., and Goodman, A. (2021) CellProfiler 4: improvements in speed, utility and usability. BMC Bioinformatics 22, 433.

11. Cai, R., Bai, P., Quan, M., Ding, Y., Wei, W., Liu, C., Yang, A., Xiong, Z., Li, G., Li, B., Deng, Y., Tian, R., Zhao, Y. G., Wu, C., and Sun, Y. (2024) Migfilin promotes autophagic flux through direct interaction with SNAP29 and Vamp8. Journal of Cell Biology 223, e202312119.

12. Buglak, D. B., Holmes, K. H. M., Galletta, B. J., and Rusan, N. M. (2024) The proximal centriole-like structure maintains nucleus–centriole architecture in sperm. Journal of Cell Science 137, jcs262311.

13. Hartley, M., Kleywegt, G. J., Patwardhan, A., Sarkans, U., Swedlow, J. R., and Brazma, A. (2022) The BioImage Archive – Building a Home for Life-Sciences Microscopy Data. Journal of Molecular Biology 434, 167505, Computation Resources for Molecular Biology.

14. Wilkinson, M. D. et al. (2016) The FAIR Guiding Principles for scientific data management and stewardship. Scientific Data 3, 160018.

15. Senft, R. A. et al. (2023) A biologist’s guide to planning and performing quantitative bioimaging experiments. PLOS Biology 21, 1–22.

16. CC BY 4.0 Deed — Attribution 4.0 International — Creative Commons. https://creativecommons.org/licenses/by/4.0.

17. Schmied, C. et al. (2024) Community-developed checklists for publishing images and image analyses. Nature Methods 21, 170–181.

18. Miura, K., and Nørrelykke, S. F. (2021) Reproducible image handling and analysis. The EMBO Journal 40, e105889.

19. Harris, C. R. et al. (2020) Array programming with NumPy. Nature 585, 357–362.

20. McKinney, W. Data Structures for Statistical Computing in Python. Proceedings of the 9th Python in Science Conference. 2010; pp 56 – 61.

21. Terpilowski, M. A. (2019) scikit-posthocs: Pairwise multiple comparison tests in Python. Journal of Open Source Software 4, 1169.

22. Virtanen, P. et al. (2020) SciPy 1.0: Fundamental Algorithms for Scientific Computing in Python. Nature Methods 17, 261–272.

23. Hunter, J. D. (2007) Matplotlib: A 2D graphics environment. Computing in Science & Engineering 9, 90–95.

24. Waskom, M. L. (2021) seaborn: statistical data visualization. Journal of Open Source Software 6, 3021. Cohen, J. *Statistical power analysis for the behavioral sciences*; Routledge, 2013.

